# Individual differences in functional connectivity during naturalistic viewing conditions

**DOI:** 10.1101/084665

**Authors:** Tamara Vanderwal, Jeffrey Eilbott, Emily S. Finn, R. Cameron Craddock, Adam Turnbull, F. Xavier Castellanos

## Abstract

Naturalistic viewing paradigms such as movies have been shown to reduce participant head motion and improve arousal during fMRI scanning relative to task-free rest, and have been used to study both functional connectivity and task-evoked BOLD-signal changes. These task-evoked changes result in cortical activity that is synchronized across subjects and involves large areas of the cortex, and it is unclear whether individual differences in functional connectivity are enhanced or diminished under such naturalistic conditions. This work first aims to characterize variability in functional connectivity (FC) across two distinct movie conditions and eyes-open rest (n=34 healthy adults, 2 scan sessions each). At the whole-brain level, we found that movies have higher intra- and inter-subject correlations in cluster-wise FC relative to rest. The anatomical distribution of inter-subject variability was similar across conditions, with higher variability occurring at the lateral prefrontal lobes and temporoparietal junctions. Second, we used an unsupervised test-retest matching (or “finger-printin”) algorithm that identifies individual subjects from within a group based on functional connectivity patterns, quantifying the accuracy of the algorithm across the three conditions. We also evaluated the impact of parcellation resolution, scan duration, and number of edges on observed inter-individual differences. The movies and resting state all enabled identification of individual subjects based on FC matrices, with accuracies between 62 and 100%. Overall, pairings involving movies outperformed rest, and the more social and faster-paced movie attained 100% accuracy. When the parcellation resolution, scan duration and number of edges used were increased, accuracies improved across conditions, and the pattern of movies>rest was preserved. These results suggest that using dynamic stimuli such as movies enhances the detection of FC patterns that are distinct at the individual level.

**Highlights:**

- Intra- and inter-subject FC correlations are compared across rest and movies.
- Movies outperformed rest in an unsupervised identification algorithm based on FC.
- Movies outperform rest regardless of parcellation, scan length, or number of edges.
- Watching movies enhances the detection of individual differences in FC.

## 1. Introduction

As psychiatric research has shifted towards a dimensional conceptualization of symptoms and behaviors (Insel et al., 2010), interest within neuroimaging has expanded to include brain-based characterization at the individual level (Arbabshirani et al., 2013). Despite the reliability of functional connectivity (FC) patterns across individuals and across testing sessions (Yeo et al., 2011; Zuo et al., 2010; Damoiseaux et al., 2006; Shehzad et al., 2009), FC relationships have also been shown to capture significant inter-individual variability, generating optimism for their eventual use as biomarkers of mental illness (Finn et al., 2015). Recent work has begun to characterize the spatial and state-based aspects of individual differences in FC.

### 1.1 Spatial aspects of FC variability.

Functional neuroimaging data sets containing retest scans have been leveraged to investigate inter-individual variability in FC patterns while controlling for intra-individual variability. Mueller et al. demonstrated that individual differences in FC were largest in association cortex including lateral prefrontal regions and the temporoparietal junction (Mueller et al., 2013). Unsurprisingly, unimodal sensory and motor regions were the least variable across subjects. At the network level, frontoparietal and ventral attention networks exhibited the largest variability in FC, followed next by the default and dorsal attention networks. This pattern of results was subsequently confirmed independently (Chen et al., 2015).

A second wave of studies extended these findings by using unsupervised test-retest sorting algorithms to match pairs of FC matrices. Finn et al. used a cohort (n = 126) of the Q2 Human Connectome Project data, and showed that correlations between FC matrices could be used to identify individual subjects from within a group (Finn et al., 2015). Further, the FC edges that contributed most to successful matching were located in the frontoparietal network. Airan et al. used a similar approach on four publicly available data sets with different sequence parameters and acquisition durations (Airan et al., 2016). They found that the brain regions which contributed most to individual subject characterization involved default, attention, and executive control networks. Taken together, these data reinforce the view that FC of heteromodal cortex is particularly variable, and importantly, that group-level variability contains differences that are distinct at the individual subject level.

Investigations of FC variability inherently capture structural variability, including differences in anatomical features such as cytoarchitecture, sulcal depth, and morphology, as well as variability in functional mapping of corresponding areas across subjects (Brett et al., 2002; Frost and Goebel, 2012; Zilles and Amunts, 2013; Shah et al., 2016). The significance of this heterogeneity is evident in the improvement attained by the “hyperalignment” of functional data (in representational space) using functional correlations rather than structural features to align data across subjects (Conroy et al., 2013; Guntupalli et al., 2016; Langs et al., 2015). At the individual level, gradients in FC profiles have been shown to delineate functionally distinct cortical areas that in and of themselves demonstrate individual-specific variation (Xu et al., 2016; Laumann et al., 2015), and methods are being developed to create individualized maps of FC relationships (Wang et al., 2015). Recent work related four areal properties (architecture, function, connectivity, and topography) to create a multi-modal parcellation of the cortex. These data showed that even with improved inter-subject alignment, specific regions demonstrated atypical topological arrangements in some subjects (Glasser et al., 2016). Though it is not yet possible to delineate the precise contribution of these underlying structural factors to inter-individual differences in FC measures, the emerging literature indicates that spatial variability is a major factor. Accordingly, multiple groups have shown that using finer-grained parcellation schemes for FC studies enhances inter-individual variability (Airan et al., 2016; Finn et al., 2015; O’Connor et al., ????). The fact that spatial variability is likely not the only factor is indicated by the presence of differences in FC variability that are present across states or conditions (see 1.2 below) and data showing that changing spatial parameters such as the degree of smoothing does not significantly alter the performance of test-retest matching algorithms (Finn et al., 2015).

### 1.2 Collection states and FC variability.

The effects of acquisition conditions on FC continue to be examined and debated (Mennes et al., 2013; Cole et al., 2014; Arbabshirani et al., 2013). A general question in the current context is whether inter-individual differences in FC are more robust under less constrained states such as rest versus tasks. As part of a thorough investigation of reliability and reproducibility in FC measures using a large database (n = 476), Shah and colleagues showed that individual patterns in FC were preserved across multiple task and rest conditions (Shah et al., 2016). Geerligs et al. investigated “state and trait” components of FC across rest and an audiovisual finger-tapping task (Geerligs et al., 2015). By comparing correlations across conditions, they showed that state and trait effects each explained approximately equal amounts of variance in FC at a single scanning session. Interestingly, when examining the temporal variability of FC, Elton and Gao found that a Stroop-like letter-naming task decreased variability relative to rest (Elton and Gao, 2015). Finn and colleagues showed that when using FC-based identification (i.e., matching) algorithm, the maximal accuracy (94%) was attained when using rest-rest correlations; accuracy decreased to 54-87% when using rest-task or task-task correlations, suggesting that individual differences are more pronounced during less constrained states, but are still present in task-based FC data. These studies indicate that inter-individual differences in FC are not abolished when using tasks, at least when the tasks are conventional and discrete such were used in these studies.

Different results have been demonstrated when using more naturalistic tasks. One study investigated inter-individual differences during movie-watching using a Hitchcock film (*Bang! You’re Dead*) (Geerligs et al., 2015). This study showed that the least amount of overlap and the highest amount of variance occurred within the movie-task comparison relative to both the movie-rest and task-rest comparisons, suggesting that perhaps movies have a unique effect on FC patterns. These data indicate that individual differences in FC are more pronounced when conditions are more constrained and engaging, or perhaps more accurately, when more networks are recruited simultaneously in a naturalistic fashion. However, to date, it remains unclear which collection states might be most advantageous for the study of FC patterns that are distinct at the individual level.

### 1.3 Movies and FC variability.

Airan et al. point out that to optimize individual subject characterization, investigators should seek to maximize inter-subject variability while minimizing intra-subject variability (Airan et al., 2016). The same concept was outlined by Wang et al. who assert that to be clinically useful, a mapping technology must demonstrate (among other things) high reproducibility within subjects while also being sensitive to functional differences between subjects (Wang et al., 2015). These are testable constructs, as outlined by Strother et al., who advocate for cross-condition comparisons of data-analytic parameters, including both prediction accuracy across data sets and reproducibility within the same data set (Strother et al., 2002). Due to the significant improvement in compliance regarding head movement and arousal levels conferred by movie watching in the scanner (Vanderwal et al., 2015), we wanted to investigate the effects of movie watching on these aspects of FC variability.

In the present study, we investigated individual differences in FC during two distinct movie watching conditions and eyes-open rest. Movie conditions were an abstract, nonverbal movie called *Inscapes* that was created to maximize compliance while minimizing cognitive load, and a verbal, social, complex clip from an action movie (*Ocean’s Eleven*, Warner Brothers 2001, directed by Steven Soderbergh). These two movies have previously been shown to elicit different FC patterns, particularly involving the default and frontoparietal networks, with Inscapes demonstrating mean FC that was more comparable to rest than to the action movie clip (Vanderwal et al., 2015). The study is divided into two parts. First, we provide an overview of aspects that relate to FC variability, including cross-condition comparisons of FC, measures of inter- and intra-subject correlations in FC, and the spatial distribution of inter-individual variability of FC. Based on these cross-condition characterizations of variability, we predicted that movies would enhance the ability to detect individual differences in FC that are distinct at the individual level. The second part of the study tests this hypothesis. We ran an unsupervised test-retest matching algorithm that used FC matrices to identify individual subjects from among a group. We also ran the algorithm using different parcellation schemes, acquisition durations, and percent of edges used to test whether these factors differentially affected the two types of movies and rest. Primary outcomes were the different accuracy percentages of the identification algorithm across the three conditions.

## 2. Materials and Methods

### 2.1 Data collection. Participants.

Healthy right-handed adults were recruited from the community, and 46 participants completed two testing sessions with a one-week interval. Twelve participants self-reported falling asleep during one or both sessions, and were excluded from further analysis, leaving n = 34 (18 females, mean age 24.4 ± 5.1 years). Data from a subset (n = 22) were published previously (Vanderwal et al., 2015). Exclusion criteria included neurological or psychiatric diagnoses, use of centrally acting medications, heavy alcohol use, any illicit drug use in the past 6 months, cardiovascular disease, significant visual or hearing impairment, and self-reporting less than six hours of sleep per night. All participants gave written consent and were compensated for their participation. The study was approved by the Human Investigations Committee at Yale University School of Medicine.

#### Procedure

Imaging was performed on a Siemens Trio 3-Tesla scanner with a 32-channel head coil. Standard structural images used an MP-RAGE sequence (TR=1900ms, TE=2.52ms, TI=900ms, flip angle=9°) yielding 1mm3 voxel size. Functional data were collected using a single shot echo planar imaging sequence (TR=2500ms, TE=30ms, flip angle=80°, voxel size=3mm isotropic) across 38 slices in the same plane as the anatomical scans. All participants completed 3 functional scans during which stimuli were presented via E-Prime software, version 2.0 (Psychology Software Tools, Pittsburgh, PA). Images were back-projected onto a screen that participants viewed via a mirror mounted on the head coil. Sound-reducing headphones over protective earplugs enabled participants to hear the soundtracks. Three 7 minute and 20 second conditions included *Inscapes* (detailed description of this movie is provided in Vanderwal et al. 2015), a clip from the movie *Ocean’s Eleven* (Warner Brothers, 2001, directed by Steven Soderbergh) referred to here as Oceans, and Rest (see Figure 1). The order of conditions was counter-balanced across participants. Each condition started and ended with 10 seconds of fixation; the first 10 seconds were dropped for all analyses. Participants were asked to watch the screen and to stay as still as possible during each condition. Foam wedges were fitted around the participant’s head for comfort and to decrease movement. Retest sessions occurred 1 week later at the same time slot whenever possible. Due to scheduling issues, 6 participants had different time slots for scan 1 and scan 2, but the 1-week interval was maintained.

**Figure 1.**
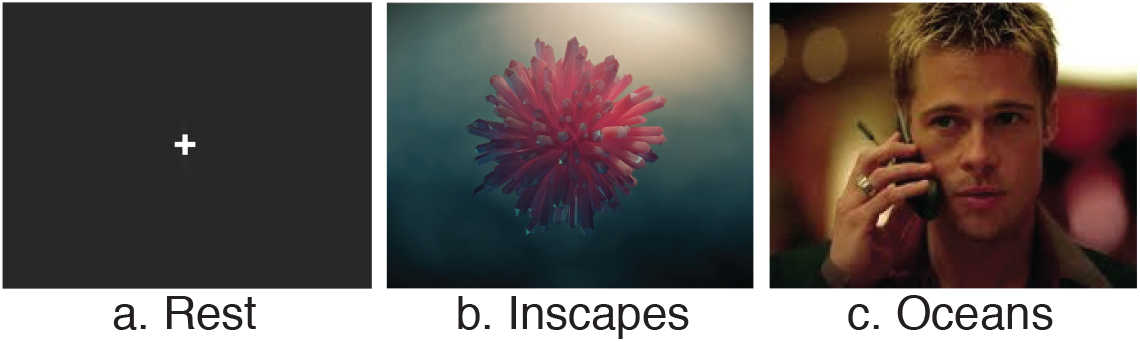
Scenes from three conditions. Conditions were a) eyes-open Rest, using a static fixation cross on a dark grey background; b) Inscapes, a nonverbal, nonsocial, abstract animation designed to maintain engagement while minimizing cognitive load; and c) a clip from vault scene of the action movie *Ocean’s Eleven*. All conditions were 7 minutes long, with 10 seconds of fixation at the beginning and end. Inscapes can be viewed and downloaded at headspacestudios.org.

#### Data processing

Standard data preprocessing was performed using the Configurable Pipeline for Analysis of Connectomes (C-PAC) including motion realignment and transformation into Montreal Neurological Institute (MNI) space using Advanced Normalization Tools (Avants et al., 2008). Nuisance signal regression removed linear and quadratic trends, motion estimates, and COMPCOR with 5 principal components (Behzadi et al., 2007), and was followed by temporal filtering (0.008-0.1Hz). Motion was evaluated using framewise displacement (FD) which quantifies head motion between each volume of functional data (Power et al., 2012). Following Finn et al., data were not spatially smoothed prior to averaging within our cluster-based regions of interest (ROIs). Number of volumes per condition was 172.

#### Whole-brain FC matrices

All subsequent analyses were based on FC connectivity matrices. Matrices were constructed using a functional parcellation scheme comprising 200 ROIs (Crad-200; Craddock et al., 2012). For each subject, we extracted the mean time series of each ROI and then computed the Pearson’s correlation coefficient between all ROI pairs to produce a 200x200 whole-brain connectivity matrix for each subject for each condition. Subsequent analyses used only unique ROI pairs (i.e., A-B and not B-A), leaving 19900 edges. Correlation coefficients were Fisher z-transformed, averaged across subjects, and then reverted to r-values to produce group-level correlation matrices. To qualitatively assess similarities across conditions in terms of the correlations within and between large-scale functional networks, we arranged the ROIs on the matrix according to network membership using the 7-network scheme (visual, somatomotor, dorsal attention, ventral attention, limbic, frontoparietal and default networks; see Figure 2), defined by Yeo and Krienen and colleagues (Yeo et al., 2011).

**Figure 2.**
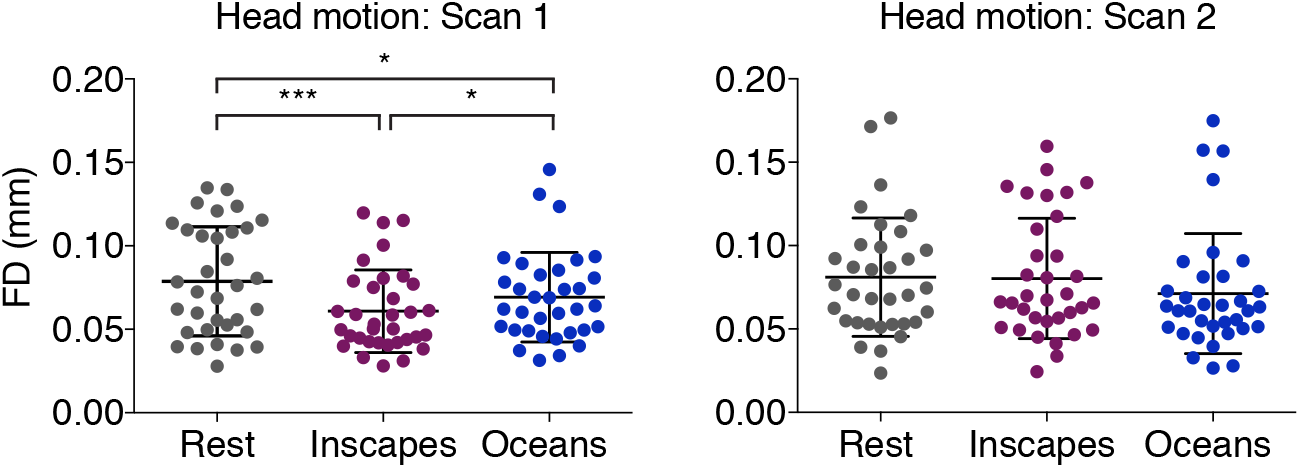
Head motion during movies and Rest (n=34, healthy adults). Head motion was evaluated by quantifying the framewise displacement (FD) between each volume of functional data (Power et al., 2012). Significant differences were found across all conditions at Scan 1, with Rest *>* Oceans *>* Inscapes. At Scan 2 (one week later), no significant differences were found in mean head movement across conditions. Overall, head motion during all acquisition conditions across both scanning sessions was low, reflecting the high compliance of this young, healthy adult population. Bracketed lines indicate which comparisons showed significant differences (*=p<0.05, ***=p<0.001).

### 2.2 FC variability across conditions

Following previous methods used to assess similarity and difference in FC matrices across conditions or states (Cole et al., 2014; Geerligs et al., 2015), we created connectivity matrices for each condition by averaging the connectivity matrices across participants within each condition. These matrices are used to visualize the group-level connectivity for movies and Rest. For statistical comparison across conditions, however, we calculated pairwise Pearson’s correlations between connectivity matrices of the different conditions (i.e. Rest-Inscapes, Inscapes-Oceans, Oceans-Rest) within a subject. To better match the noise estimate (see below), we used half the volumes by computing the Pearson’s correlation coefficient between the first half of condition 1 and the first half condition 2, as well as the second half of condition 1 and condition 2. The average of these two correlation coefficients was squared, and this squared value represents the proportion of variance that is similar or shared across those conditions. It is assumed that the remaining proportion of variance represents across-condition variance, such that:

- r = correlation between two conditions
- r^2^ = shared variance between those conditions, termed overlap
- (1 - r^2^) = total remaining variance, assumed to include both state-based and noise-based contributions

To estimate the noise contribution to this between-state variance, we calculated the split-half correlation within each condition. The split-half correlation obtained for Rest was used to estimate noise regardless of the pair of conditions being compared, as movies are not expected to be consistent from the first half to the second half.

- r_*sh*_ = split-half correlation of Rest
- (1 - r_*sh*_^2^) = estimate of variance due to noise

This facilitates the subtraction of noise from the total variance, such that:

- (1 - r^2^) - (1 - r_*sh*_^2^) = state-based variance

It is important to note two differences between our method and that outlined in Geerligs et al. (2015). First, all computations were performed using pairwise Pearson’s correlations between connectivity matrices of the different conditions *within* a subject, so we expect lower cross-condition correlations overall. Second, as explained above, the between-condition correlation coefficients are based on only half of the volumes, again likely returning lower r-values than has been shown previously.

#### Intra- and inter-subject FC correlations

Intra-subject (or within subject) correlations of FC were computed by first calculating the Pearson’s correlation coefficient of the two scanning sessions’ FC matrices for each subject. To compute the inter-subject correlations, we performed the same procedure between one subject and every other subject within a single scanning session. The pairwise correlations were averaged to provide a single inter- and intra-subject r-value for each subject.

#### Spatial distribution of inter-individual variability

Following an approach outlined in Mueller et al., we wanted to map cluster-level inter-subject variability of FC using a method that accounted for intra-subject variability (Mueller et al., 2013). First, group-level inter-subject variance values were obtained as follows: for each condition’s first scanning session we computed a correlation coefficient for each ROI (across all 199 of its edges) between all possible subject pairings. The resulting r-values were squared and subtracted from 1 to convert them to measures of dissimilar variance. Averaging across subject pairs then yielded an estimate of total inter-subject variance at each cluster. Next we computed the intra-subject variance for each condition by performing the same procedure, this time correlating between each subject’s scan 1 and scan 2 FC matrices, yielding a 200 x 34 (clusters x subjects) matrix, which was subsequently averaged across individuals to obtain a group-level map. Using ordinary least-squares regression, the intra-subject variance was regressed out of the total inter-subject variance, and the residuals were taken to represent the inter-subject variability. We did not regress out a measure of technical noise. Residual values were mapped onto surface space using CARET (Van Essen et al., 2001). To assess the variability by network, we used the 7-network schema from Yeo and Krienen et al., averaging the variability across all clusters belonging to each network (Yeo et al., 2011).

### 2.3 Accuracies of identification algorithm

The prediction procedure closely followed methods described elsewhere (Finn et al., 2015). In brief, six databases were created, one for each of the three conditions for both Scans 1 and 2. Each database consisted of the Crad-200 FC matrix for each subject for a given condition (34 matrices per data set). To run the matching algorithm, two databases were selected at a time. A subject’s FC matrix was selected from one, and the Pearson’s correlation coefficient was then calculated between that matrix and every matrix in the other database. The two matrices with the highest correlation were deemed the “matched pair,” and the accuracy of the algorithm was simply the percentage of correct pairs when checked against the known subject identities. We ran the algorithm across testing sessions and across conditions, resulting in a structure of 30 pairings (e.g., Rest 2-Rest 1, Oceans 2-Rest 2, Oceans 2-Rest 1). Because of our moderate sample size, we wanted to be sure that accuracies did not reflect chance pairings. We thus performed nonparametric permutation testing in which false identity pairs were randomly assigned and the algorithm was run 1000 times to determine how many times the false pair was identified as being the most strongly correlated. To investigate the role of head motion in the matching, we computed discrete motion distribution vectors for each participant based on the framewise displacement time courses across all 3 conditions and across both scanning sessions. The mean and standard deviations of the FD across all subjects and conditions was computed, and 60 bins were set to capture the grand mean +/- 3 s.d. and vectors were calculated accordingly. The 1x60 vectors were then used in the same way that the FC matrices were to run the identification algorithm. This procedure tests whether each individual’s motion characteristics can be used to identify individuals from within a group, and helps to assess the degree to which motion might contribute to the FC-based matching algorithm.

#### Parcellation resolution

To test if the resolution of the parcellation had differential effects on identification accuracy across conditions, we parcellated the data at all of the 43 resolutions defined in a publicly available atlas that used a spatially constrained spectral clustering approach of independent resting state data (Craddock et al., 2012). The range of the number of clusters was 10-950. We then ran the identification algorithm using each parcellation on the Scan 2-Scan 1 within-condition pairings. All subsequent analyses used the Crad-200 parcellation and only the Scan 2-Scan 1 within-condition pairings.

#### Scan duration

To test if shorter scan durations handicapped the matching accuracy of one condition more than another, we ran the same matching algorithm, varying the amount of data used between two volumes and the full 172 volume run, starting from the beginning of the run and adding sequential TRs one at a time.

#### Number of edges

To test if one of the conditions required fewer edges in order to make the correct identity matches, we sequentially tested the algorithm using increasing numbers of edges. To dictate the order in which we added edges, we calculated the differential power (DP) for each edge (Finn et al., 2015). DP provides a measure of how contributory an edge is to successful matching. Values indicate the proportion of the time a subject is matched to itself rather than to another subject based on that edge. We then rank-ordered the edges from least contributory (lowest DP) to most contributory (highest DP) within each condition. Next, we ran the matching algorithm using only the lowest 0.5% of edges, and successively repeated this procedure adding an additional 0.5% at each increment until 100% of the edges were used.

## 3. Results

### 3.1 Compliance

Subjects self-reported falling asleep during 14 Rest runs, 7 Inscapes runs, and zero Oceans runs. The remaining 34 subjects had little overall head motion, with mean FD=0.07mm (s.d.=0.03) at scan 1, and a mean FD=0.08mm (s.d.=0.04) at scan 2 (see Figure 2). At the first scanning session, mean FD was significantly lower for both movies relative to Rest, and Inscapes was significantly lower than either comparison condition (one-way repeated measures ANOVA, F_(2,33)_=9.589, p=0.0004, post hoc two-tailed t-test, Inscapes-Rest p=0.0001, Oceans-Rest p=0.05, Inscapes-Oceans p=0.019). At the second scanning session, no significant differences in head motion were found (one-way repeated measures ANOVA, F_(2,33)_=1.855, p=0.17).

### 3.2 Characterizing FC variability

Within-condition split-half correlations for FC matrices were as follows: Rest=0.64, Inscapes=0.65, Oceans=0.60 (see Figure 3). There was a significant effect of condition on split-half correlation (one-way repeated measures ANOVA, F_(2,33)_=7.202, p=0.0015). Follow-up paired t-tests showed no significant difference in the split-half correlations between Inscapes and Rest (t_(33)_=0.411, p=0.68), but Oceans was significantly different from both Inscapes (t_(33)_=3.581, p=0.0011) and Rest (t_(33)_=3.149, p=0.0035). Cross-condition comparisons using half of the volumes showed moderate correlations, with Rest-Inscapes r=0.54, Inscapes-Oceans r=0.48 and Oceans-Rest r=0.47 (one-way repeated measures ANOVA, F_(2,33)_=26.08, p<0.0001, post hoc two-tailed t-test, Inscapes-Rest vs. Inscapes-Oceans p<0.0001, Rest-Inscapes vs. Rest-Oceans p<0.0001, and Oceans-Rest vs. Oceans-Inscapes p<0.041. These r-values are lower than those reported by Geerligs et al. for similar comparisons, possibly because we maintained within-subject pairings and used only half of the volumes when calculating the cross-condition correlations. Rest and Inscapes had the highest overlap at 29% and the lowest state-based difference (after subtracting out a noise estimate) of 12.9%.

**Figure 3.**
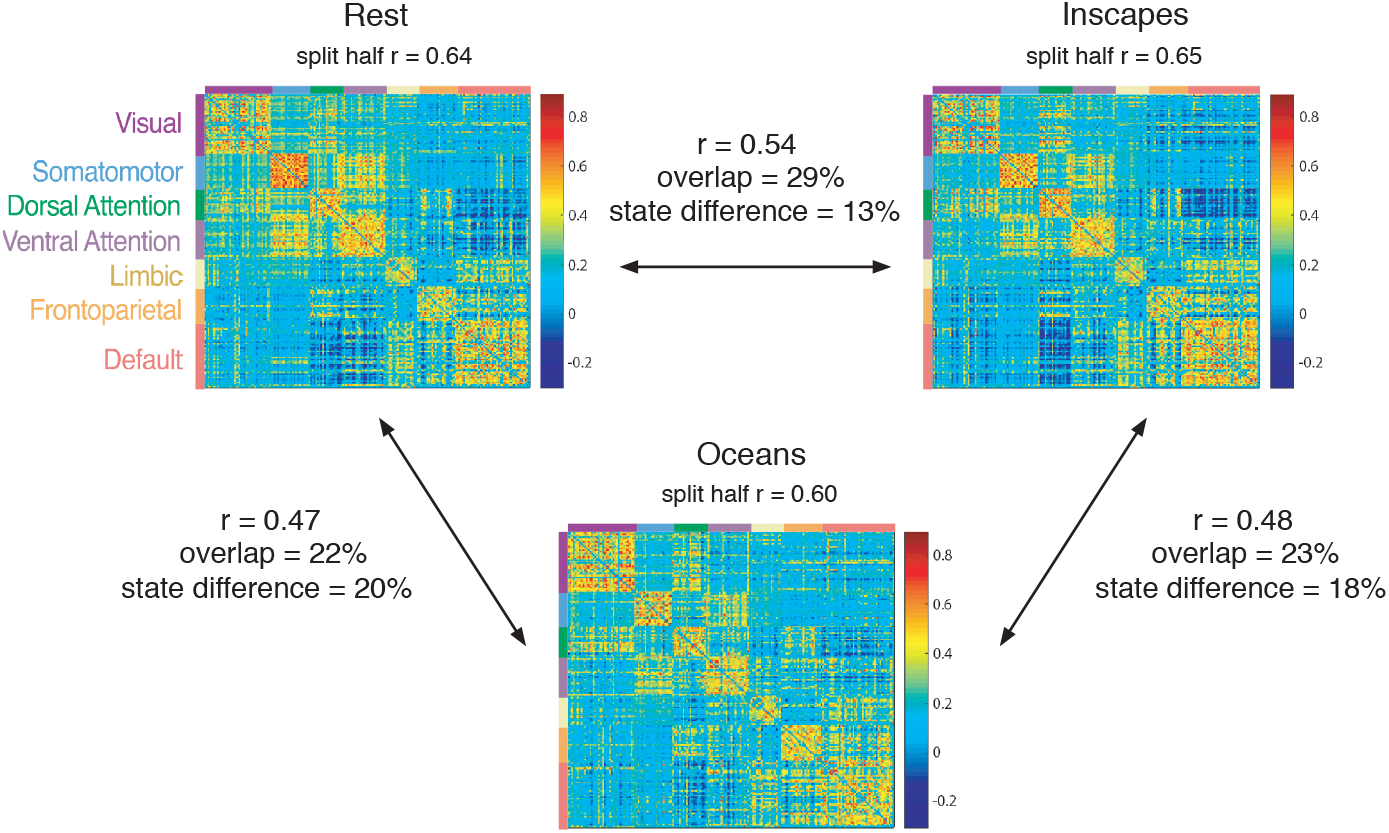
Group-level similarity and variance in FC matrices across movies and Rest (n=34, healthy adults). Pearson’s correlation coefficients were calculated between each pair of conditions to produce the r-value denoted in the matrices. Rest and Inscapes are the most strongly correlated conditions with the highest amount of overlap and the lowest state difference. Rest and Inscapes also have the highest split-half correlations. These data align with our previous report suggesting Inscapes is associated with FC patterns that more closely resemble Rest than those of conventional movies (Vanderwal et al., 2015).

#### Intra- and inter-subject FC correlations

Intra-subject correlations for FC were uniformly stronger than inter-subject correlations (see Figure 4). For intra-subject correlations, movies were stronger than Rest, but were not different from each other (one-way repeated measures ANOVA, F_(2,33)_=4.29, p=0.019, post hoc two-tailed t-tests, Inscapes-Rest p=0.01, Oceans-Rest p=0.017, Oceans-Inscapes p=0.82). For inter-subject correlations, movies were again stronger than Rest, and were again not significantly different from each other (one-way repeated measures ANOVA, F_(2,33)_ =30.34, p<0.0001, post hoc two-tailed t-tests, Inscapes-Rest p<0.0001, Oceans-Rest p<0.0001, Oceans-Inscapes p=0.17).

**Figure 4.**
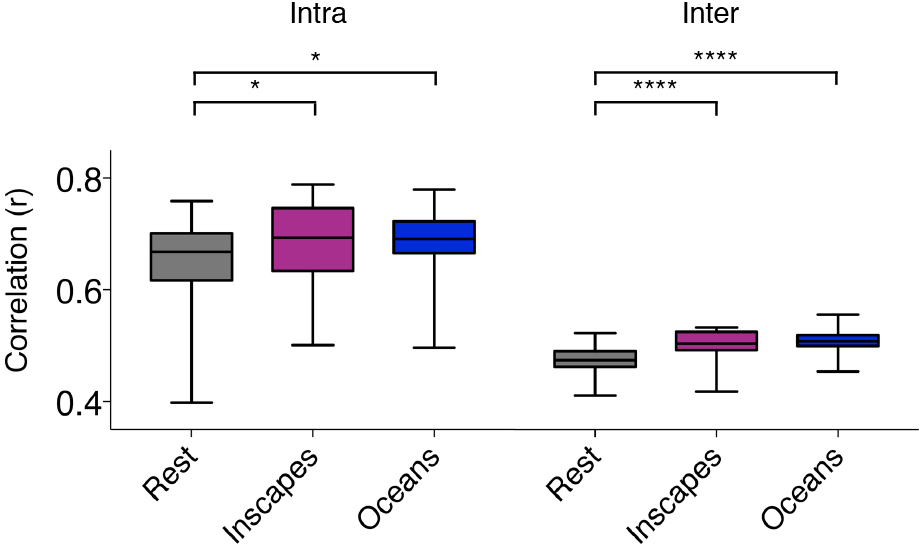
Intra- and inter-subject FC correlations. Intra-subject correlations of whole-brain, cluster-based FC were greater than inter-subject correlations for all conditions. Both movies had significantly greater intra-subject correlations relative to Rest, with no significant difference between movies. The same pattern was found for inter-subject correlations, with movies greater than Rest (*=p<0.05, ****=p<0.0001).

#### Spatial distribution of FC variability

Inter-subject FC variability (by cluster) demonstrated a nonuniform spatial distribution with higher variability in the lateral prefrontal lobes, temporoparietal junctions, and along regions of the lateral temporal lobes (see Figure 5). Lower variability was found in primary sensory and motor cortices. This pattern is similar to previous reports (Mueller et al., 2013) and was similar across both movies and Rest. Inscapes had higher variability in temporal regions, while Oceans had higher variability in prefrontal regions. When calculated for each of 7 networks, FC variability was highest in the frontoparietal network and lowest in the visual and somatomotor networks across conditions. Within the frontoparietal network, variability was highest for Oceans.

**Figure 5.**
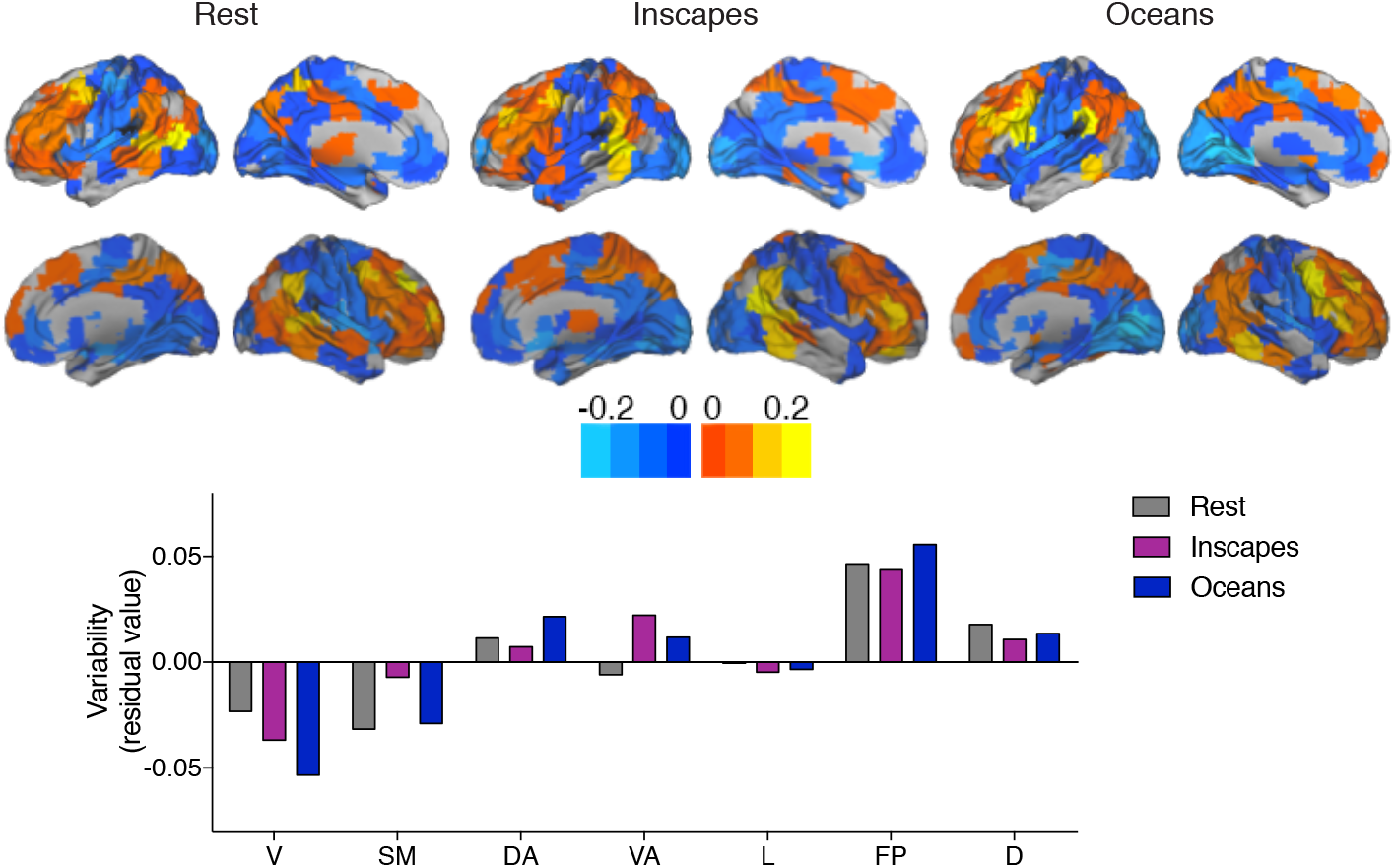
Spatial distribution of inter-individual variability in FC. Inter-individual variability was quantified for FC of each cluster after least-squares regression to correct for underlying intra-subject variability. Consistent with previous work (Mueller et al., 2013), FC variability is lowest in primary sensory and motor regions, and highest in heteromodal regions such as lateral prefrontal cortices and the temporoparietal junctions. Qualitatively, this pattern of spatial distribution occurs across all three scanning conditions. Variability by network was highest in the frontoparietal network and lowest in visual and somatomotor networks. V=visual, SM=somatomotor, DA=dorsal attention, VA=ventral attention, L=limbic, FP=frontoparietal, D=default, masks for networks based on Yeo and Krienen et al., 2011.

### 3.3 Accuracies of identification algorithm

We first tested prediction accuracy using the Crad-200 parcellation, and found high accuracies across and within conditions with a range of 62-100% (see Figure 6). Oceans attained 100% accuracy, and in general, the highest accuracies were associated with pairings that included movies. Importantly, high accuracies were attained for cross-condition pairings, indicating that individually distinct patterns in FC persisted across conditions. The permutation testing (performed 1000 times to quantify the percentage of time the algorithm matched a randomly assigned “false identity” pairing) had an average accuracy of 0.36% (with maximal accuracy of 5.9%) for Scan 2 to Scan 1 pairings, and 0.51% (with maximal accuracy of 8.8%) for Scan 1 to Scan 2 pairings. When using motion distribution to match subjects, accuracies ranged from 9-24%. Results for permutation testing and motion distribution are shown in Supplementary Materials.

**Figure 6.**
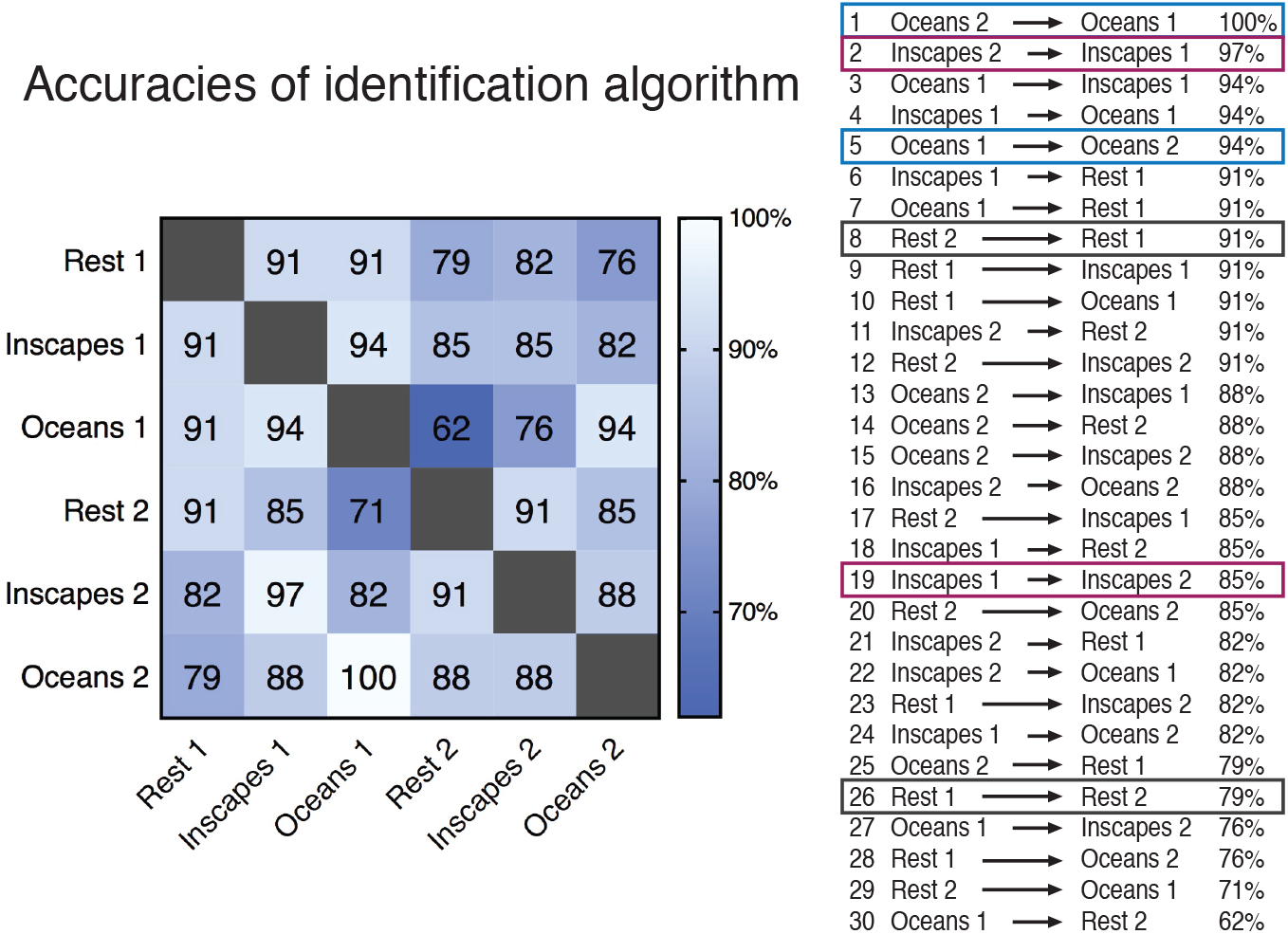
Accuracies of unsupervised test-retest matching algorithm based on FC matrices (n=34, healthy adults). Individual subjects were correctly identified by the unsupervised algorithm across all conditions, with accuracies ranging from 62-100%. Overall, the highest accuracies were associated with pairings that involved movies. High accuracies were attained in cross-condition matches, indicating that individual subjects have distinct FC patterns that persist across conditions.

#### Parcellation resolution

When tested at different parcellation resolutions, the within-condition Scan 2-Scan 1 identification accuracies improved with higher resolutions, as expected (Figure 7a). Varying the resolution had differential effects by condition: Oceans attained 100% accuracy at 120 clusters, Inscapes at 500 clusters, and Rest hit a ceiling accuracy of 97% at 650 clusters.

**Figure 7.**
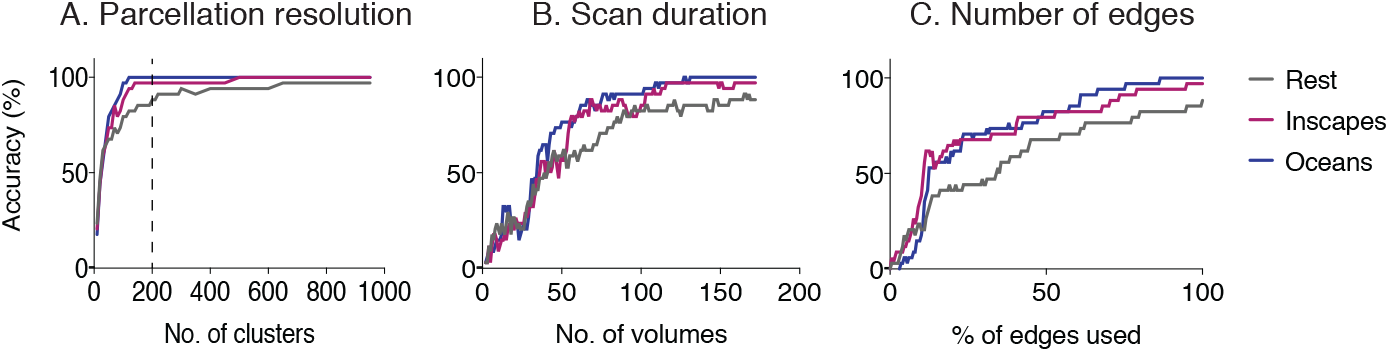
The effect of varying data parameters on accuracy of unsupervised matching algorithm (n=34). **A. Parcellation resolution:** Using the within-condition matching algorithm (Scan 2-Scan 1) on data parcellated at different resolutions, accuracies increased with higher resolutions. Oceans attained 100% accuracy at 120 clusters, Inscapes at 500 clusters, and Rest hit a ceiling accuracy of 97% at 650 clusters. Dashed line highlights the Crad-200 matrix, which was used for all subsequent analyses. **B. Scan duration:** Including more volumes led to an increase in accuracies for all conditions. Oceans reached 100% accuracy at 127 volumes; Inscapes reached 97% at 116 volumes; and Rest hit its ceiling of 91% at 165 volumes. **C. Number of edges used.** After rank-ordering edges according to differential power (DP), we sequentially added edges that were increasingly contributory to successful matching. The more edges used, the higher the accuracy across conditions. Both movies had similar accuracies until about 75% of the edges were used, at which point Oceans diverged from Inscapes. Overall, varying each of these parameters did not alter the general pattern of movies outperforming Rest with regards to the matching of individual subjects.

#### Scan duration

Also as expected, longer scan durations positively affected the accuracy of the algorithm. Oceans reached 100% accuracy at 127 volumes, Inscapes reached maximal accuracy of 97% at 116 volumes, and Rest reached a ceiling accuracy of 91% at 165 volumes (Figure 7b).

#### Number of edges used

Again, including higher numbers of edges produced higher prediction accuracies across all conditions, as was expected. Oceans and Inscapes demonstrated almost overlapping graphs up until 75% of the edges were included, at which point Oceans had higher accuracies than Inscapes. Overall, movies outperformed Rest at all numbers of edges included (Figure 7c).

## 4. Discussion

### 4.1 Accuracies of identification algorithm for movies and rest

This study investigated the effects of movie watching on individual differences in FC. We showed that an unsupervised test-retest matching algorithm that identifies subjects from within a group based on FC performed well using data acquired during both movies and Rest, and that the highest accuracies were attained using movies.

Overall accuracies of the matching algorithm ranged from 62-100%. These results are in-line with data from Finn et al. who reported accuracies of 54-94% in a larger sample using rest and task (Finn et al., 2015). The highest accuracy in our data (100%) was attained when matching FC matrices between scan sessions of Oceans, with Inscapes reaching 97%, and Rest 91%. The pattern of these accuracy relationships (Oceans *>* Inscapes *>* Rest) held true across parcellation resolutions, scan durations, and number of edges used. We conclude that relative to task-free resting state conditions, movie watching preserves— and possibly enhances—the ability to detect differences in FC patterns that are distinct at the individual level.

### 4.2 Variability in FC of different acquisition conditions

The matching algorithm used is based on correlations of FC matrices between separate scanning sessions. Consequently, we would expect intra-subject correlations to play a substantial role in the success of the algorithm. When examining cluster-wise, whole-brain FC, our data showed that movies had significantly stronger intra- and inter-subject correlations relative to Rest. Additionally, intra-subject correlations were stronger than inter-subject correlations for all conditions, in accordance with the success of the matching algorithm across conditions.

Further, previous work has shown that FC edges which contributed most to successful identification matches were found in the frontoparietal network (Finn et al., 2015). In our data, variability within the frontoparietal network was greatest for Oceans, perhaps contributing to the observed 100% accuracy attained for the Oceans-Oceans pairings. In general, the spatial distribution of inter-subject variability in FC during Rest and movies followed the same pattern as had been previously reported during Rest: the lowest variability occurred in primary motor and sensory cortices with higher variability in heteromodal cortex involving the prefrontal and temporal cortices (Mueller et al., 2013; Chen et al., 2015). These data suggest that the spatial distribution of inter-individual variability observed during Rest is not drastically shifted by movie watching, and more specifically, that variability in the frontoparietal network is high across conditions.

### 4.3 Individually distinct FC and naturalistic paradigms

To date, the majority of studies utilizing naturalistic paradigms have focused on the concerted nature of BOLD-signal changes evoked by movie watching that have been shown to involve large areas of the cortex (Hasson et al., 2004, 2010; Kauppi et al., 2010). Intuitively, one might assume that because of this shared activation across subjects, patterns of FC would be less distinctive at the individual level. Our data indicate that this is not the case as the highest accuracies of the matching algorithm were attained using movie-watching data. In addition to the high intra-subject FC correlations discussed above, another possible contributing factor to this pattern is that concerted activity across subjects in multiple voxels enables individually distinct patterns of FC to “stand out” more. In other words, we hypothesize that movie watching may not itself evoke individual differences in functional neural responses, but that the whole-brain processing at the group-level that occurs during naturalistic paradigms enhances the detection of individually distinct FC patterns. This seems plausible given the fact that cross-condition pairings in our data also attained high accuracies, indicating that the same individually distinct patterns are maintained across conditions.

Relatedly, Papageorgiou et al. suggested that complex stimuli might elicit useful shifts in whole-brain signal-to-noise ratios when they used a highly engaging task in which subjects received real-time feedback to their neural responses during a silent counting task (Papageorgiou et al., 2013). They posited that frontoparietal regions and the insula regulated global processes during engaging conditions, conferring an improvement in signal-to-noise ratios. When taken together with data indicating that individual variability is highest in frontoparietal networks (Mueller et al., 2013), and that matching algorithms rely heavily on edges contained in the frontoparietal network (Finn et al., 2015), we suggest that during naturalistic conditions, the frontoparietal network may play a dual role in the identification of individually distinct FC patterns. First, frontoparietal FC itself may comprise individually distinct differences, and second, frontoparietal control may cause an advantageous shift in broader processes enhancing the detection of individually distinct FC patterns. Whatever the mechanism, studies to date, including the data presented here, indicate that the frontoparietal network plays a key role in the detection of individual differences in FC.

### 4.4 Compliance

The major compliance advantage of using movies in healthy adult populations relates to arousal levels. In this study, subjects self-reported falling asleep during 14 Rest runs, 7 Inscapes runs, and zero runs of Oceans. In line with our previous report, head movement at the first scan was significantly better during both movies relative to rest. A new finding reported here is that Inscapes showed significantly lower head movement than Oceans. However, no differences in head movement were found at the second session. We speculate that habituation and loss of novelty may have contributed to this null finding.

Because head motion has previously been shown to be more trait-like than state-like (Siegel et al., 2016; Couvy-Duchesne et al., 2014), we were concerned that head motion might contribute to the accuracy of the matching algorithm. When using motion distribution as the basis for an identity algorithm (i.e., with no FC measures), accuracies ranged from 6-24%, which is higher than chance but much lower than the accuracies attained using FC measures (62-100%). We conclude that motion likely contributes to the matching of FC matrices, but that it does not account for the primary finding that naturalistic conditions enable the detection of FC differences at the individual level.

### 4.5 Limitations and future directions

The study design did not include a rich phenotypic assessment, and consequently, we were not able to test for correlations between FC variability during movie watching and clinically relevant behaviors or traits. Because movies appear to enhance the ability to detect individual differences in FC, some brain-behavior relationships may be identified using naturalistic paradigms that are not detectable using conventional tasks. For example, a pediatric study was able to identify math-based brain-behavior relationships using fMRI data collected during Sesame Street clips that were not detected using a well-validated conventional fMRI math task (Cantlon and Li, 2013). Given the absence of cross-condition differences in head motion at the second scanning session, further work examining head movement patterns would be helpful to better understand the utility of movies with regard to compliance for repeated measures. Other groups have begun to look at individual differences in temporal dynamics of FC during task reorganization (Chai et al., 2016; Simony et al., 2016), and it would be interesting to investigate these relationships under the ongoing dynamic stimulation provided by naturalistic conditions. Finally, future studies that incorporate multi-modal or group-weighted parcellation schema (Glasser et al., 2016; Mejia et al., 2016) or explorations of FC in non-anatomical space (Conroy et al., 2013) might be combined with the use of naturalistic viewing paradigms to further enhance the sensitivity of fcMRI to identify individually distinct patterns of FC.

### 4.6. Conclusions.

1. Movies preserve, and possibly enhance, the ability to detect patterns in FC that are distinct at the individual level.
2. Movies had stronger intra- and inter-subject correlations in FC relative to Rest, and inter-individual variability in the frontoparietal network was highest during Oceans. These factors may facilitate the observed 100% matching accuracy attained using Oceans.
3. Compliance benefits of using movies with healthy adults center around arousal levels, with half as many subjects self-reporting sleep during Inscapes relative to Rest, and no subjects self-reporting sleep during Oceans. Significant head motion advantages for healthy adults occurred, but only at the first exposure to the stimuli.
4. Movies may be advantageous for future efforts to identify brain-behavior correlations in pediatric and psychiatric populations.

---

**Supplemantary Table 1.** Permutation testing of unsupervised identification algorithm with falsely assigned identity pairs. The algorithm was run 1000 times using false pairings. The mean accuracy attained for Scan 2-Scan 1 pairings was 0.36%, with a highest accuracy of 5.9%. For Scan 1-Scan 2 pairings, the mean accuracy was 0.51%, with a highest accuracy of 8.8%. The table shows the number of times the algorithm identified the false pairing as being the most correlated pairing. These data indicate that correct pairings are identified by the algorithm at rates that far exceed chance (62-100%), lending validity to the high percentages attained when correctly identifying subjects.

| Number of<br>times correct<br>(out of 1000) | Scan 1 -><br>Scan 2 | Scan 2 -><br>Scan 1 |
| --- | --- | --- |
| 0 | 857 | 866 |
| 1 | 137 | 127 |
| 2 | 5 | 6 |
| 3 | 1 | 1 |

**Supplemantary figure 1.**
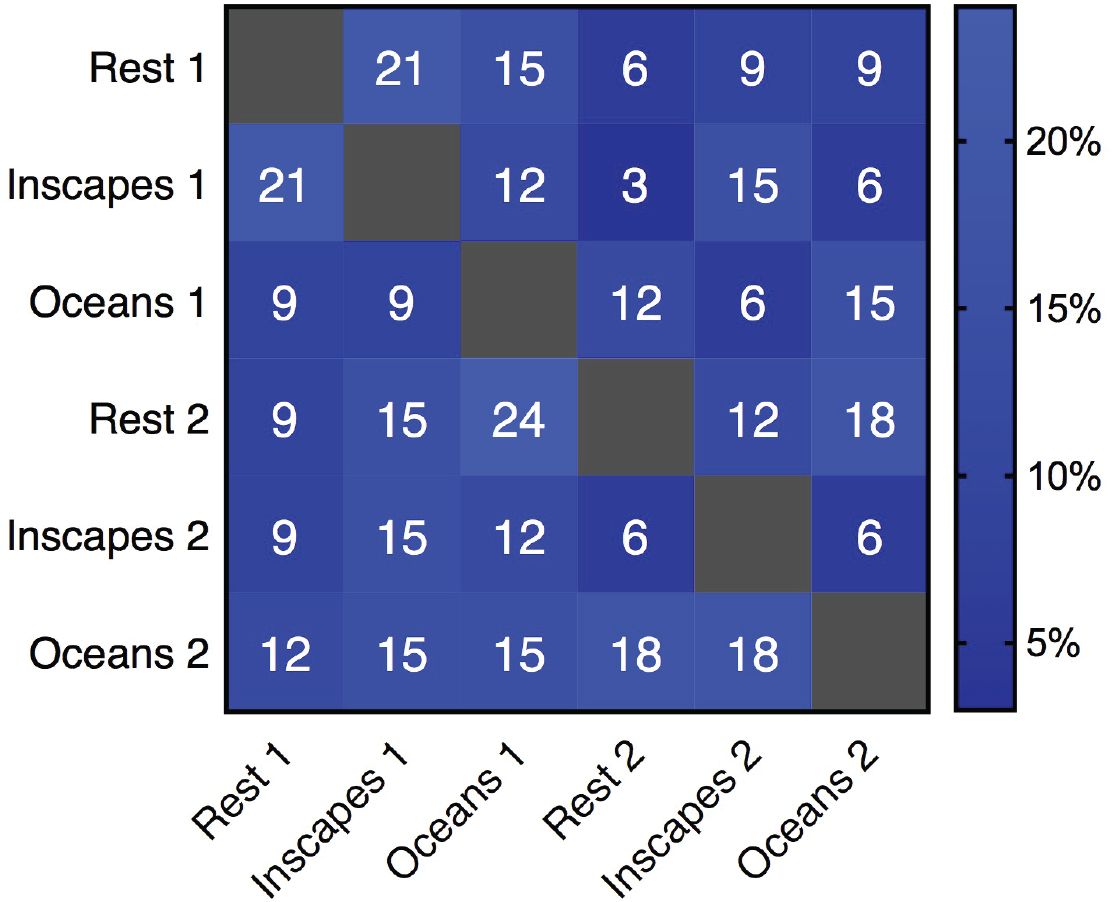
Accuracies of matching algorithm based on motion distribution. Accuracies attained using only motion parameters ranged from 6-24%. These numbers are greater than chance, and likely reflect the fact that motion distribution is a trait that is somewhat identifiable across individuals. However, these accuracies are much lower than those attained using FC matrices (62-100%). Motion likely contributes to the FC matching, but it does not appear to account for the main effect.

